# Hair glucocorticoids are not a historical marker of stress – exploring the time-scale of corticosterone incorporation into hairs in a rat model

**DOI:** 10.1101/2020.03.27.012377

**Authors:** Pernille Colding-Jørgensen, Sara Hestehave, Klas S.P. Abelson, Otto Kalliokoski

**Author notes:** Correspondence to: Otto Kalliokoski, Department of Experimental Medicine, Blegdamsvej 3B (Panum building, office 16.1.34), DK-2200, Copenhagen N.

## Abstract

Hair glucocorticoids are increasingly popular biomarkers, used across numerous research fields as a measure of stress. Although they are suggested to be a proxy of the average HPA axis activity spanning a period of weeks or months into the past, this theory has never been tested.

In the present study, adrenalectomized rats with no endogenous (adrenal) glucocorticoid production were used to study how circulating glucocorticoid levels would be reflected in the glucocorticoid levels found in hair samples. By dosing the animals daily with high levels of corticosterone for seven days, while sampling hairs before, during, and after treatments, a timeline for glucocorticoid uptake into hairs was constructed. This kinetic profile was compared to two hypothetical models, and the theory that hair glucocorticoids are a record of historical stress had to be rejected.

Corticosterone concentrations in hairs were found to increase within three hours of the first injection, the highest concentrations were found on the seventh day of treatments, and the decrease in concentrations post-treatment suggests rapid elimination. We speculate that hair glucocorticoid levels can only be used to characterize a stress-response for a few days following a postulated stressor.

An updated model, where glucocorticoids diffuse into, along, and out of hairs needs to be adopted to reconcile the experimentally obtained data. The inescapable consequence of this updated model is that hair glucocorticoids become a marker of – and can only be used to study – recent, or ongoing, stress, as opposed to historical events, weeks or months in the past.

## 1. Introduction

### 1.1 Background

Measuring hair glucocorticoids (hGCs) as biomarkers of stress in humans and other animals has gained momentum over the past decade. The popularity of quantifying hGCs is due to a number of factors: Hairs are sampled non-invasively, are stored easily (at room temperature), and can be processed through relatively simple means^1^. However, the greatest reason for quantifying glucocorticoids (GCs) in hairs, as opposed to other biological matrices, has to do with time-scale.

Within minutes following a stressful event, GC levels in the bloodstream will increase as the body manages its acute stress response^2^. GCs permeate all tissues and the increase is quickly mirrored in secretions such as saliva^3^, tears^4^, and sweat^5^. As the body discards excess GCs, increases will also be seen in urine^6^ and fecal matter^7^. Depending on the species of study, the stress-associated increase in GCs can consequently be measured in excreta hours to days after a major stress response. This is something that has been used to great effect in wildlife biology, where studying stress in animals is common, but where it is not desirable to catch and restrain the species of study^8,9^. The example was set in a landmark study by Creel, Creel, and Monfort who could demonstrate, using only non-invasive means – by sampling feces – that dominant African wild dogs and dwarf mongooses appeared to experience greater levels of social stress than their lower ranking conspecifics^10^. Because excreted GCs provide an average of circulating GC levels over several hours – for some species, days^11^ – these methods have been shown to be useful across fields of study. Take humans for example. GC levels in the bloodstream will fluctuate wildly and increase for brief periods in response to momentary exertions^12^, both physical and mental – running to catch a train; searching for misplaced car keys; speaking in front of a large audience. It is often of interest to separate these momentary stressors from the chronic stress of, for example, working night shifts^13^ or suffering from a chronic pain condition^14^. For these situations, GC levels in urine samples are useful as they approximate the total circulating GCs integrated between voidings^15^, as opposed to the easily-confounded spot-checks offered by blood sampling. GC levels in hair are suggested to be an even more powerful tool, as they purportedly collect GC levels from the bloodstream for the entirety of the hair’s growth^16,17^. This premise allows for studies into even more subtle long-term stressors, such as stress brought on by a constrained diet in wildlife^18^ or the bio-energetic stress brought on by habitual substance abuse in humans^19^.

A tool that returns the average HPA-axis activity over a period of several months already has great utility, but, in addition, quantitating hGCs is suggested to offer something unique. As the growing hair purportedly retains traces of periods of stress, the hair could potentially be read (lengthwise) like the rings of a tree or a core sample of Antarctic ice. Each segment of a hair can be traced to a point in the past where it grew out of the follicle, and the GC content of that segment is suggested to act as a time capsule, retaining a memory of the HPA axis activity at the time^20,21^. A single strand of hair would thus present a “stress calendar”, collecting a record of past stress reaching months into the past^22^. Embracing this theory, archeologists have reconstructed whole segments of the lives of long dead mummies, from an allostatic point-of-view, by analyzing the GC content in locks of thousand-year-old hair^23^. With powerhouse studies such as these, it is easy to understand the method’s popularity. Adoption of hGC analyses has happened with great speed across several disciplines: whether used to assess mental health in humans^24,25^, welfare in farm animals^26,27^, or distress in laboratory animals^28,29^ (to mention just a few applications).

There is one major issue however: In the rush to fulfil the potential of the method, rigorous validation has been eschewed. The means by which GCs enter the hair are currently based on guesswork: It is assumed that the contribution of GCs in sweat^30^ and sebum^31^ are negligible to hGC levels^32^; it is believed that production of GCs in the hair follicle^33^ does not confound measurements^34^; it is hoped that localized release of GCs^35^ does not ruin studies. Moreover, the molecular interactions making the hair retain GCs are all but unknown^36–38^. An elucidation of the kinetics of GC incorporation into hairs is long overdue.

### 1.2 Theory

The present investigation sought to compare experimentally obtained data to two hypothetical models of hGC kinetics. A scenario was crafted where the two models predicted radically different results. This allows for simple assessment of theory, allowing one or both of the models to be disproven.

The popular model (outlined in *e.g*., Russell et al.^16^, Meyer & Novak^39^, Stalder & Kirschbaum^40^, Wester & van Rossum^41^, and Hodes et al.^42^) proposes that glucocorticoids are 1) deposited into the growing hair at the level of the follicle; 2) that they remain largely stationary in a particular segment as the hair grows; 3) that limited post-deposition degradation of glucocorticoids happens. The unknowns of the model are often emphasized: other routes for incorporation are, for example, occasionally discussed^43–45^ and treatments of hair in humans (dyeing, shampooing, *etc*.) has been noted to leach out glucocorticoids^46–49^. However, these three pillars are necessary to be able to, for example, subsection hairs, inferring stress in periods in the past. Whether made explicit, or simply implied, we feel that this is an accurate representation of the theory underlying a majority of hair glucocorticoid investigations.

Conversely, it has been noted that most, if not all, published results of glucocorticoid analyses can be explained under an alternate frame of interpretation^38^. This alternative model proposes that glucocorticoids are 1) deposited at the level of the follicle; 2) travel freely along the shaft of the hair; 3) are degraded, eliminated, or equilibrate with surrounding tissue, to a degree where the hair cannot retain its concentration of glucocorticoids.

The difference between the two models is that the former – we will call this the Classic model (borrowing the terminology of Grass *et al*.^32^) – allows for hGCs to be a long-term marker of (chronic) stress. Under this paradigm, GCs measured in a hair summarize the total HPA axis activity for a period reaching weeks, if not months, into the past. Additionally, it allows for subsectioning and utilizing hairs as the aforementioned stress calendar. The latter hypothesis – we will call this the Alternate model – suggests that hair glucocorticoids are not a historical marker of stress at all. Under this paradigm, hGCs are only useful for assessing the concurrent stress – a summary of HPA axis activity collecting hours, or at most days, of recent past. Moreover, under this model the idea of a “stress calendar” is impossible.

### 1.3 Hypotheses

By utilizing adrenalectomized (ADX) rats with no adrenal GC production, a controlled stressor of considerable magnitude can be simulated through repeated injections of corticosterone (CORT), the principal effector GC in rats^8^. By sampling hair before, during, and after the period of injections (chosen to be seven days), using a shave-reshave protocol (the principle originally established by Davenport et al.^50^), a coarse model of hGC kinetics can be established (Figure 1). The GC concentrations in shaved hairs can be compared, in turn, to the time-concentration profile of hGCs predicted by the two hypothetical models (Figure 2).

**Figure 1.**
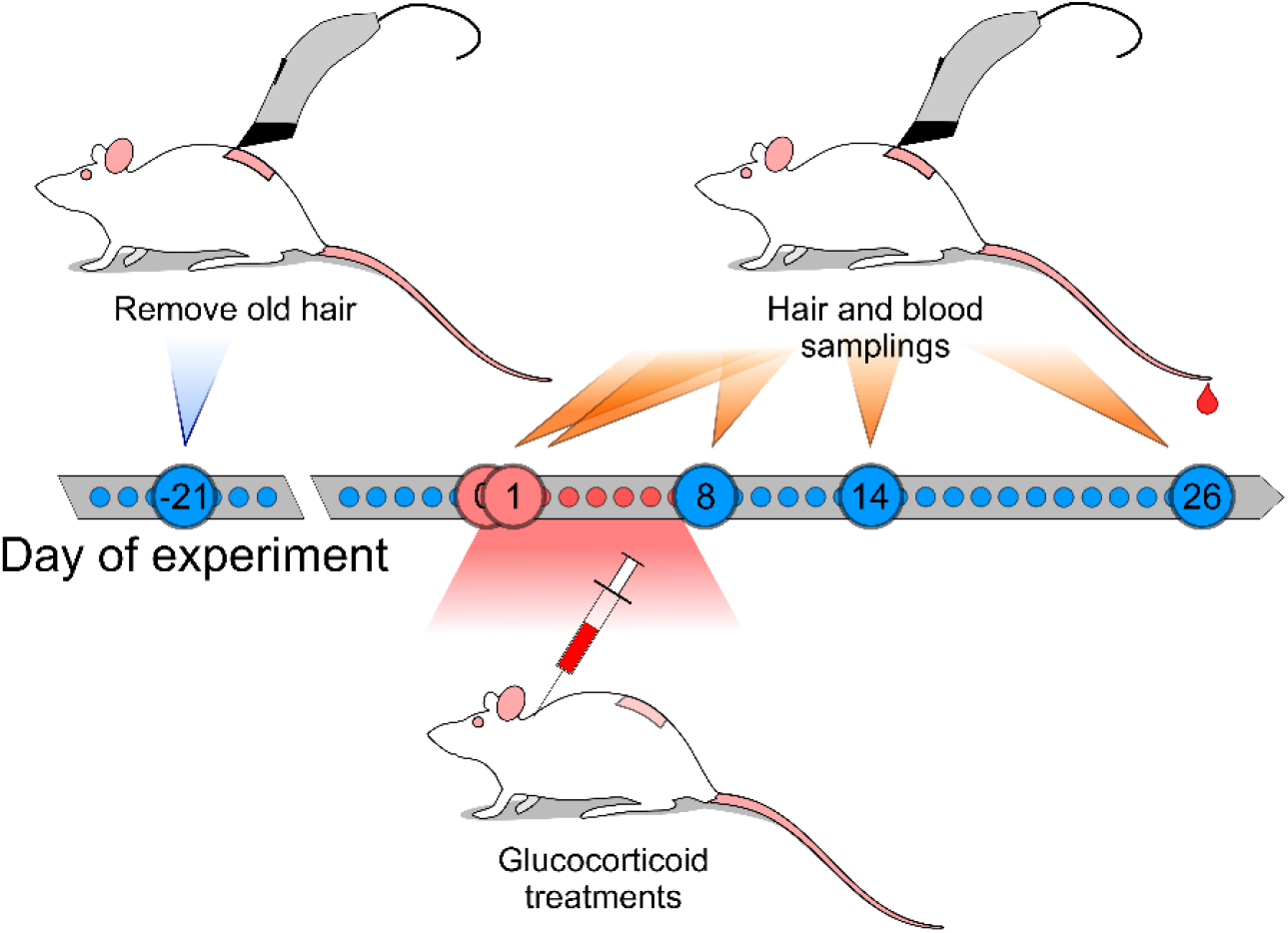
The design of Experiment 2 or the present study. Employing a shave-reshave protocol using adrenalectomized (ADX) rats treated with corticosterone, we can investigate the kinetics of hGC. Because of the time required to regrow enough hair needed for analysis, the same rat can unfortunately not be sampled at each time point. Instead, a larger cohort has to be studied.

**Figure 2.**
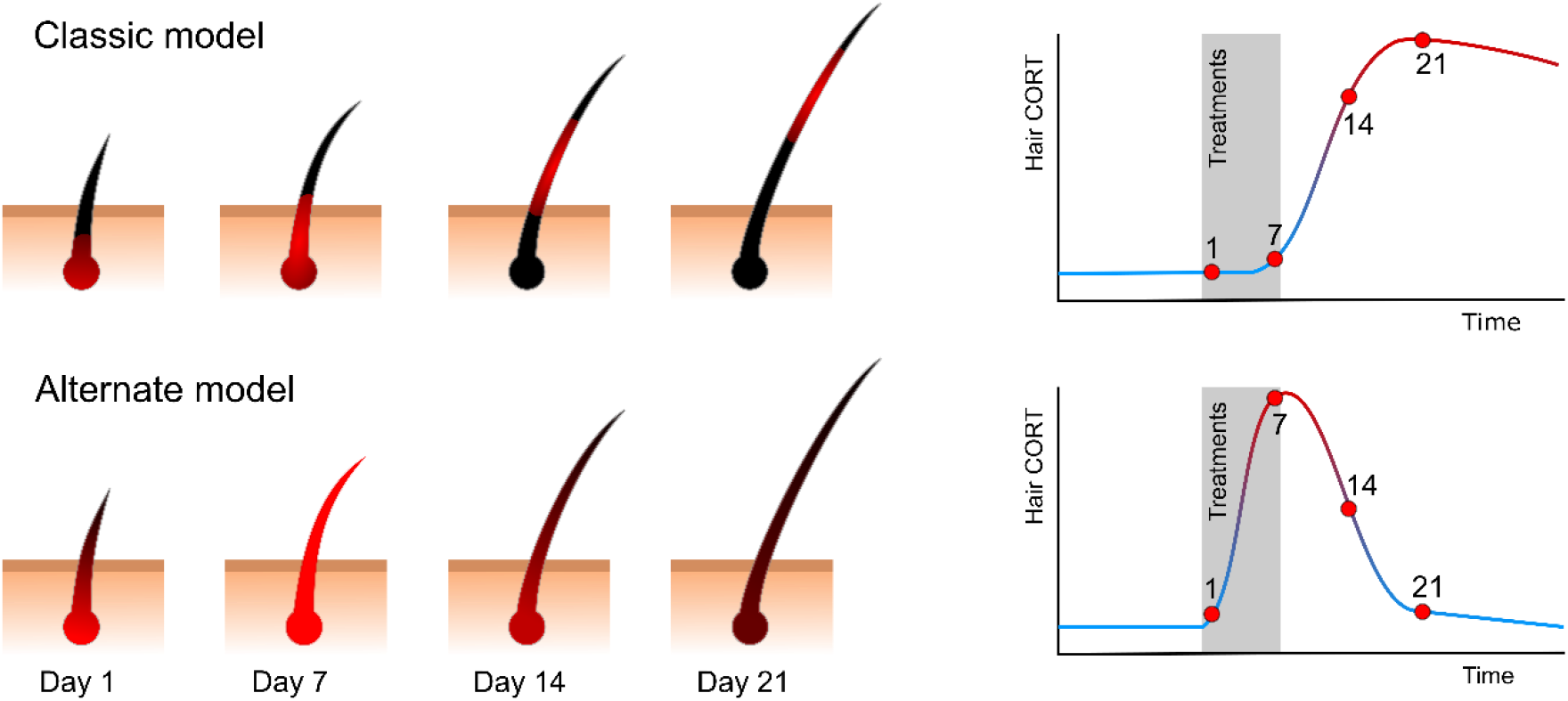
Hypothetical (simplified) scenarios based on two competing models. Sampling only the part of the hair that protrudes over the surface of the skin (by shaving), we expected the measured concentrations of CORT in samples to change over time in a predictable fashion. The time-concentration profiles would look distinctly different however. The Classic model predicted a delayed CORT response to the treatments, a slow continued increase, also after treatments are terminated, and retained high concentrations for a long time. The Alternate model, by contrast, predicted much faster kinetics with a fast rise to peak CORT concentrations, a rapid decline ending in a return to baseline levels. By comparing the real-life concentration-time profiles to the two hypothetical figures, we expected to be able to reject one, or both, of the outlined models.

Specifically, three points of divergence were of interest when examining the data obtained in the present study:

1. The Classic model predicts a lag time between initiation of treatments and the consequent increase in hGC levels, since only the part of the hair protruding through the skin will be sampled when shaving. Because it takes time for hair to form in the follicle, and for it to emerge above skin, it has been suggested that an increase in hGC levels cannot be seen for several days after the onset of a stressor (the lag time has been suggested to be as long as 1-2 weeks^16,40^). By contrast, the Alternate model suggests that GCs taken up by the hair at the level of the follicle can, through longitudinal transport, be recovered in the parts of the hair that have already emerged through the skin with minimal delay.
2. An additional consequence of the proposed lag time is that the Classic model predicts a continued increase in hGC concentrations even as the stressor subsides, as hair grown under more stressful conditions continues to emerge above the surface of the skin. The Alternate model instead suggests a rapid decline with peak hGC concentrations seen during – not after – the artificial stressor.
3. Finally, under the Classic model, the hair retains its GC content for a long time (weeks to months). The hGC concentration will principally only be reduced by new GC-depleted hair emerging. This will reduce the overall hGC concentration (expressed per gram of hair), essentially “diluting” the signal. The Alternate model suggests, instead, that GCs will diffuse rapidly out of the hair, hGC concentrations eventually returning to pre-stressor levels (for ADX animals – practically GC-free hairs).

We hypothesized that the Alternate model would better describe real-life data. Consequently, we hypothesized that: 1) a measurable increase in CORT would be found in hairs within 24 h of the first injection; 2) a reduction – not a continued increase – in CORT levels would be seen in hairs obtained more than 24 h following the last injection; 3) hair CORT levels would return to baseline levels within two weeks of the final injection.

## 2. Material and methods

The study was split into two experiments. An initial experiment was conducted to test that treatments with CORT would give a measurable response in hairs. This was to serve as a basis for continued testing, however due to complications with the outsourced surgeries, a separate design was utilized for follow-ups (utilizing in-house surgeries). Since the animals would be slightly different in age and the exact study design slightly altered, results from the two experiments are kept separate – the initial experiment used as a pilot investigation.

### 2.1 Animals

#### Experiment 1

Male Sprague-Dawley rats (6 weeks old, approximately 225-250 g on arrival) were obtained from Janiver Labs (Le Genest-Saint-Isle, France). A subset of 28 rats had been adrenalectomized (ADX) approximately one week (> 7 days) prior to transport. The ADX rats were accompanied by 18 age-matched intact rats.

#### Experiment 2

Male Sprague-Dawley rats (n = 36), with an average body weight of approximately 200 grams (5 weeks old at the time of arrival) were obtained from Janvier Labs to undergo in-house adrenalectomies.

### 2.2 Housing conditions (*both experiments*)

All animals were housed in groups of three animals per cage. Individually ventilated cages (“NexGen 1800”, Allentown Inc., Allentown, NJ, USA) were lined with aspen chips (Tapvei, Kortteinen, Finland); equipped with cardboard tubes (Lillico, Horley, UK) and plastic inserts (“JAKO”; Molytex, Glostrup, Denmark) for shelter; nest-building material (Lillico) and bite bricks (Tapvei) were placed on the cage floor and changed as needed; feed (“Altromin 1314F”; Altromin GmbH, Lage, Germany), saline solution, and tap water were provided *ad libitum*. Ventilation was kept at 60 h^−1^ air changes on a cage level, temperature at 22 °C (± 2 °C [range]), and the humidity at 55 % (± 10 %). A diurnal rhythm was maintained with a 12:12 hour light-dark cycle with 30 minutes of artificial twilight at transitions and lights on at 06.00. Cages were changed every 7 days, transferring only shelters from the old cage. The same groupings of rats and housing conditions were kept throughout the study.

### 2.3 Study outline

#### Experiment 1

After 7 days’ acclimatization upon arrival from the breeder, a patch of hair on the rats’ backs was shaved with clippers. Hair was allowed to regrow for 21 days before a baseline hair sample was collected immediately before a baseline blood sample. Another 14 days later, daily treatments (subcutaneous injections) with 5 mg corticosterone (per animal) were initiated. On day seven of treatments, hair samples were collected from all animals, and serum samples were collected the next day.

#### Experiment 2

After 13 days’ acclimatization upon arrival from the breeder, bilateral adrenalectomies were carried out. Animals from one or two cages underwent surgery per day and all of the surgeries were performed within a period of 12 days. One week after the final surgeries, a patch of hair on the rats’ backs was shaved with clippers and blood samples were collected. Three weeks after collecting baseline serum samples, daily treatments (subcutaneous injections) for seven days with 5 mg corticosterone were carried out. Six animals were left untreated, forming a control group (for reference levels). Hair and blood samples were collected 3 hours, 24 hours, 8 days and 14 days after the first treatment (refer to Figure 1 for an illustration of the design). The final hair samples were collected 26 days after the first treatment and the animals were put down. The same subjects could, for obvious reasons, not be sampled at all time points – a minimum amount of sample material is needed for analysis and hair simply does not grow fast enough. In an effort to make the most of each subject, however, the animals sampled at 3 and 24 h, were the same subjects to be sampled on the final day of the study (at this time they had regrown enough hair to yield analyzable samples). Control animals were sampled 3 hours after initiation of treatments and on day 26.

### 2.4 Surgeries (*Experiment 2*)

The animals were anesthetized with a single subcutaneous (s.c.) injection of Hypnorm (0.20 mg/kg [bodyweight] fentanyl and 6.25 mg/kg fluanison; Skanderborg Apotek, Skanderborg, Denmark) and midazolam (3.13 mg/kg; Hameln Pharmaceuticals, Gloucester, UK). The absence of righting and paw-withdrawal reflexes were confirmed and the surgical area was shaved and washed (“Nex Clorex C2 surgical scrub”; Nex Medical Antiseptics, Casorezzo, Italy). The animal was placed on its back on a pre-heated operation pad (approx. 34 °C), covered with sterile draping, and eye ointment and a nose-cone providing 1 l/min pure oxygen were applied. The abdominal cavity was accessed through a mid-line incision (3-4 cm in length, starting at the *processus xiphoideus*) and the adrenals were located using blunt dissection. Throughout the extirpation, care was taken not to touch the gland directly, instead manipulating the adjacent tissue with forceps. The adipose tissue and blood vessels were cut using microsurgical straight scissors and hemostasis of the vessels was applied, as needed, using nonwoven swabs moistened with saline. The muscle layer and peritoneum were closed using simple continuous sutures (“Coated Vicryl 5-0”; Ethicon, Somerville, NJ, USA) and the skin layer was sutured intradermally (“Monocryl 6-0”; Ethicon). The same surgeon carried out all operations between hours 8.00-13.30. Aseptic technique was employed throughout and all instruments and materials were pre-sterilized.

### 2.5 Post-operative care (*Experiment 2*)

Immediately following the surgeries, animals were given carprofen (5 mg/kg, “Norodyl vet.”; ScanVet Animal Health, Fredensborg, Denmark), butorphanol (1 mg/kg, “Butamidor”; Salfarm, Kolding, Denmark), and 1 ml saline s.c., while still under anesthesia. For recovery, the animals were placed on a soft operating pad in their home cage. For the first hours (approximately 4 h), the cage was placed on an external heating source. On the first two days post-surgery, the animals’ standard feed was supplemented with energy-dense diet (“DietGel Boost”; ClearH2O, Portland, ME, USA) and softened feed pellets to counteract post-operative weight-loss. Post-operative pain-relief was given s.c. for the first three days after surgery: buprenorphine (0.05 mg/kg, “Vetergesic vet.”; Orion Pharma Animal Health, Espoo, Finland) given twice daily, and carprofen (5 mg/kg) given once daily (omitted on the last day). The first buprenorphine injection was given approximately four hours after completed surgery, and was combined with another 1 ml s.c. saline injection to prevent dehydration. For the remainder of the study, a bottle containing saline (9 g/l, prepared from tap water) was added to each cage, allowing the animals control of their osmotic balance in the absence of adrenal aldosterone. For the first three days after surgery, corticosterone (25 mg/l, prod. no. 27840; Sigma-Aldrich, Munich, Germany) was added to the drinking saline, aiding recovery.

### 2.6 Drop-outs

#### Experiment 1

Unexpected mortalities of ADX rats were encountered in the acclimatization period immediately after arrival from the breeders, where they had been adrenalectomized. Nine animals were euthanized or found dead during the first twelve days post-arrival (before start of experiments). The fast onset, symptoms – extreme lethargy, ataxia, sudden refusal to ingest feed and water, and no obvious signs of complication from the surgeries (confirmed through necropsies) – and timing, suggests that the mortalities were due to stress-induced adrenal crisis brought on by transport and exposure to a new environment. In addition, seven animals were excluded from the analysis due to their having regenerated adrenal tissue (as has been described by Gotlieb *et al*.^51^). Early indications of adrenal tissue regeneration came in the form of faster weight gain (ADX counteracts obesity in rats^52^), slower hair growth (GCs retard hair growth^53^ and ADX, consequently, accelerates it in rats^54^), and consumption of tap water (as opposed to saline; ADX animals will favor saline^55^). Adrenal tissue regeneration was confirmed by measuring serum corticosterone levels and, ultimately, through necropsies^51^. For detailed descriptions of excluded individuals, refer to the Supplemental Materials.

#### Experiment 2

One animal was removed from the study on the first day after surgery due to ruptured sutures (the animal was immediately euthanized). Animals that were suspected of having regenerated adrenal tissue were excluded from the study (n = 8). These rats were kept as companion animals in the cages for the remainder of the study, but their hair was not analyzed.

### 2.7 Treatments

A stable emulsion of 50 mg/ml corticosterone was prepared similar to what we have described previously^56^. The emulsion was employed as an injectable to create predictable circulating levels of CORT similarly to what has been described in the past^57,58^. Briefly, 500 mg corticosterone (27840; Sigma-Aldrich) was suspended in 1 ml 96 % ethanol and added to 10 ml sesame oil under stirring. The mixture was heated, evaporating the ethanol, and once the oil had become translucent, the heated solution was transferred to a sterile borosilicate glass. The tube was inverted until use to avoid the milky emulsion separating. For the injections, 100 μl of the emulsion was deposited subcutaneously in the nape of the neck. As some irritation was noted around the injection site, the inguinal region was injected for the final three days of treatments in Experiment 2.

### 2.8 Samplings

The subjects were restrained by hand and a 35 cm^2^ patch on the lower back was shaved using electrical clippers. A translucent stencil measuring 7 × 5 cm, overlaid the fur (lengthwise, centered over the spine), was used as a target area in Experiment 2 to standardize the method as best as possible. When re-sampled, the previously shaved area was still visibly different and could be sampled without using the stencil. Hair was collected onto a piece of paper, collected using tweezers, and placed in a filter bag made of cellulose fiber (“Telia”; Dansk Tefilter, Otterup Denmark). The filter bags were sealed into labelled plastic tubes and stored at room temperature until analysis. Blood samples were obtained immediately after shaving. The lateral tail veins were punctured at the level of the tail tip using a lancet or needle, and light pressure was applied as the tail was massaged in a distal motion, gently encouraging bleeding. Sample volumes of approximately 150 μl were collected into non-coated collection tubes, spun down within a few hours from collection, and fresh serum was frozen at −20 °C until analysis. Larger blood samples were collected on the last day of the study. Animals were swiftly concussed and decapitated; trunk blood was subsequently collected in 5 ml microcentrifuge tubes.

### 2.9 Processing of hair samples

Prior to being processed, tubes containing hair samples were randomly ordered (using a random number generator) and recoded, thus blinding the analyst, ensuring unbiased readings of analyses. Samples were processed similarly to what has been described by others previously^28,59,60^. The filter bags containing hairs (used to ensure none of the sample was lost in the process) were washed twice in isopropanol and left to dry in a fume hood overnight. Dried hairs were pulverized in a ball mill (5-10 min at 30 Hz, “Retsch MM 400”; Retsch GmbH, Haan Germany) and extracted in methanol overnight (solvent ratio: 50 ml/g). Pulverized samples of less than 10 mg were excluded from further analysis. The samples were centrifuged (20 min at 3,000 g followed by 15 min at 10,000 g) to pellet solids and clear supernatants were collected. The extracts were subsequently dried (“Genevac EZ-2 personal evaporator”; SP Industries, Warminster (PA), USA) and resuspended in PBS overnight.

### 2.10 Analyses

Blood samples and hair extracts were both analyzed using a commercially available competitive ELISA (“Corticosterone ELISA”, EIA-4164; DRG Instruments GmbH, Marburg, Germany) with which we have ample experience and which has previously been used to quantitate CORT in rat hairs^61^.

### 2.11 Ethics statement and justifying cohorts

The study was conducted in an AAALACi (Association for Assessment and Accreditation of Laboratory Animal Care International) accredited facility under the supervision of a local ethics committee. All procedures were carried out in accordance with EU directive 2010/63/EU and approved by the Animal Experiments Inspectorate under the Danish Ministry of Food, Agriculture and Fisheries (license number 2016-15-0201-01007).

We chose to prioritize the reduction principle of the 3Rs over carrying out the experiment in both sexes, which could potentially have increased our number of subjects due to increased inter-individual variance but provided very little additional information. Males were chosen to minimize the effect of cross-reacting hormones: the selected glucocorticoid assay is known to have higher cross-reactivities with progestogens than androgens (highest known cross-reaction: 7.4 % with progesterone). Group sizes for Experiment 2 were not based on a sample size calculation for a hypothesis test but, instead, on the ability to discern a general trend in data. For groups of five or more subjects, pilot data (log-transformed) from Experiment 1 suggested that the 95 % CI of the mean was less than 25 % (24.6 %) of the total magnitude of the response at peak (treatment) and less than 5 % (3.7 %) at baseline, which were considered rough target figures sufficient for producing a non-ambiguous time-concentration profile describing the kinetics of hair glucocorticoids. With a high degree of adrenal regeneration (found during Experiment 1), the study was over-dimensioned to account for drop-outs.

### 2.12 Statistics

CORT release is a pulsatile^12^, multiplicative process^62^, resulting in CORT concentrations best being described assuming log-normal distributions^63^. Consequently, log-transformed data were used for all analyses, geometric means are reported throughout, and confidence intervals assume log-normal distributions. Having decomposed the model characteristics into three testable hypotheses – refer to section 1.3 – only single-time-point comparisons were made. Welch’s t-tests (reporting t-values, degrees of freedom, and associated p-values) were used for all comparisons. The hypotheses were considered independent of one-another, consequently no alpha-level corrections have been employed.

## 3. Results

### 3.1 Experiment 1

With subjects lost to transport stress, subjects excluded due to their regenerating adrenal tissue, hair samples too small for analysis, and serum samples excluded as two animals did not receive CORT injections on the eight day of treatments, the final dataset was much too small to offer an accurate picture of the kinetics. We will for this reason refrain from stringent hypothesis testing and instead note the reasons for why the design was not expanded. The animals were relocated to an adjacent room in batches for blood samplings and we were consequently not surprised to find that the serum CORT levels for all intact animals were elevated (Figure 3; 95 % CI: 434-754 nM, n = 28). Similar elevated levels were found in CORT-treated ADX animals (95 % CI: 133-1,730 nM, n = 5).

**Figure 3.**
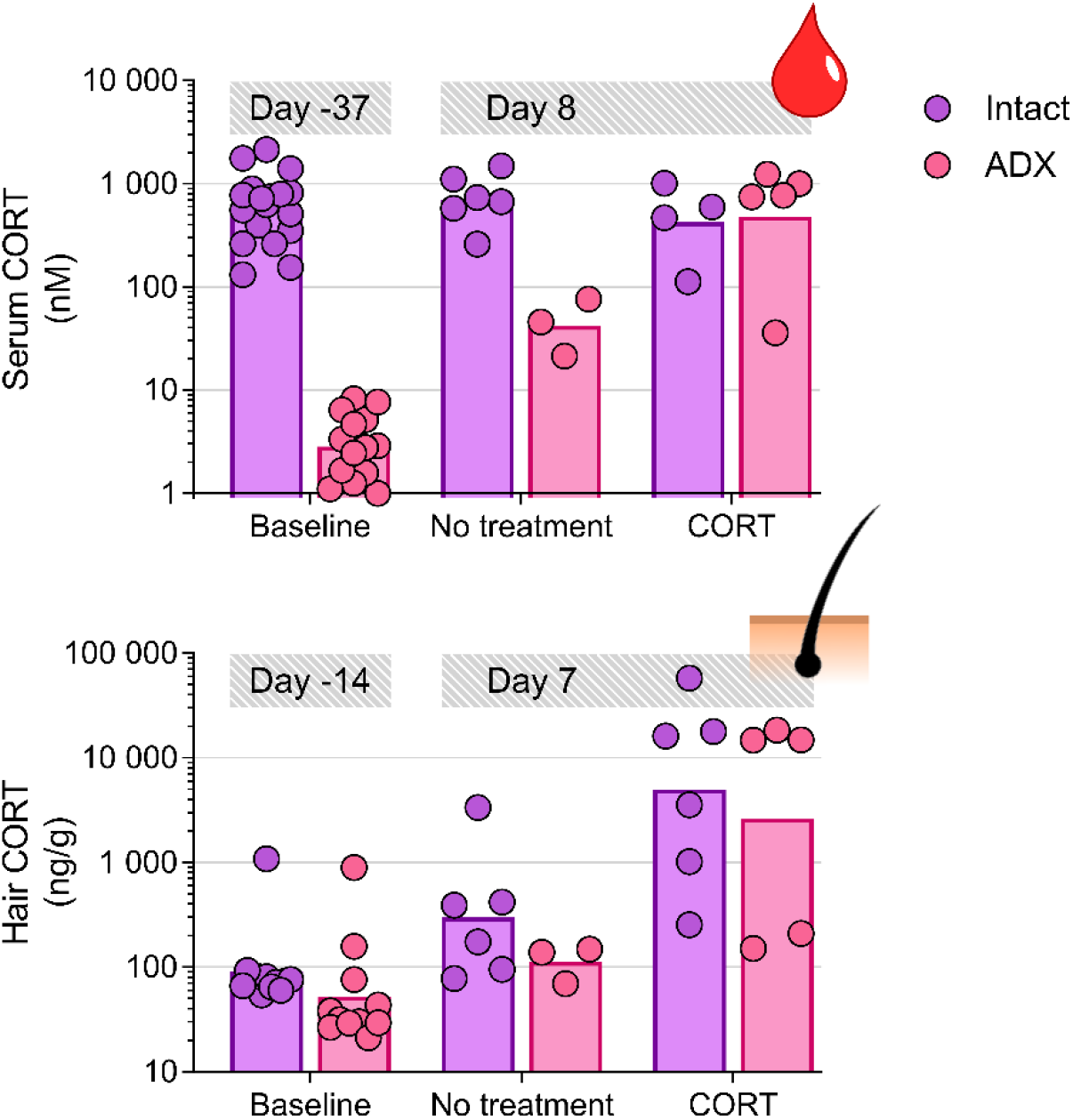
Time-concentration curves for serum and hair CORT in relation to one week’s treatments. Intact and ADX animals had similar levels of hair CORT throughout. We were however unable to obtain undisturbed blood samples from intact animals. We have refrained from hypothesis testing of data because of the underpowered design.

Hair CORT levels appeared to be similarly elevated for both ADX animals as well as intact animals from daily CORT treatments. On the seventh day of treatments the levels were nearly two orders of magnitude higher than baseline hair CORT levels (pooled estimates: Baseline – 95 % CI: 45.7-106 ng/g, n = 23; Day 7 (CORT) – 95 % CI: 885-15,741 ng/g, n = 11). There appeared, however, to be an increase in both circulating serum CORT and hair CORT for untreated subjects over time, also for the ADX animals. This suggested a regeneration of adrenal tissue in ADX animals, also in animals where no patches of regenerated adrenal tissue could be macroscopically identified during necropsy.

### 3.2 Experiment 2

Carrying out surgeries in-house, and establishing *a priori* exclusion criteria, offered a much more controlled study design and conclusive findings. Only one subject had to be removed after sampling and analysis. Serum CORT levels of control animals remained low throughout (95 % CI: 24-38 nM, n = 5) and can be explained as the assay cross-reacting with other steroid hormones (Figure 4). Treatments with CORT resulted in circulating serum somewhat higher than those seen in the pilot study (95 % CI: 3,080-4,560 nM, n = 11). 24 hours after cessation of CORT treatments, serum levels were still elevated relative to baseline (95 % CI: 498-1,270 nM, n = 5). This is likely due to the oil-based vehicle used for the injections providing a depot effect.

**Figure 4.**
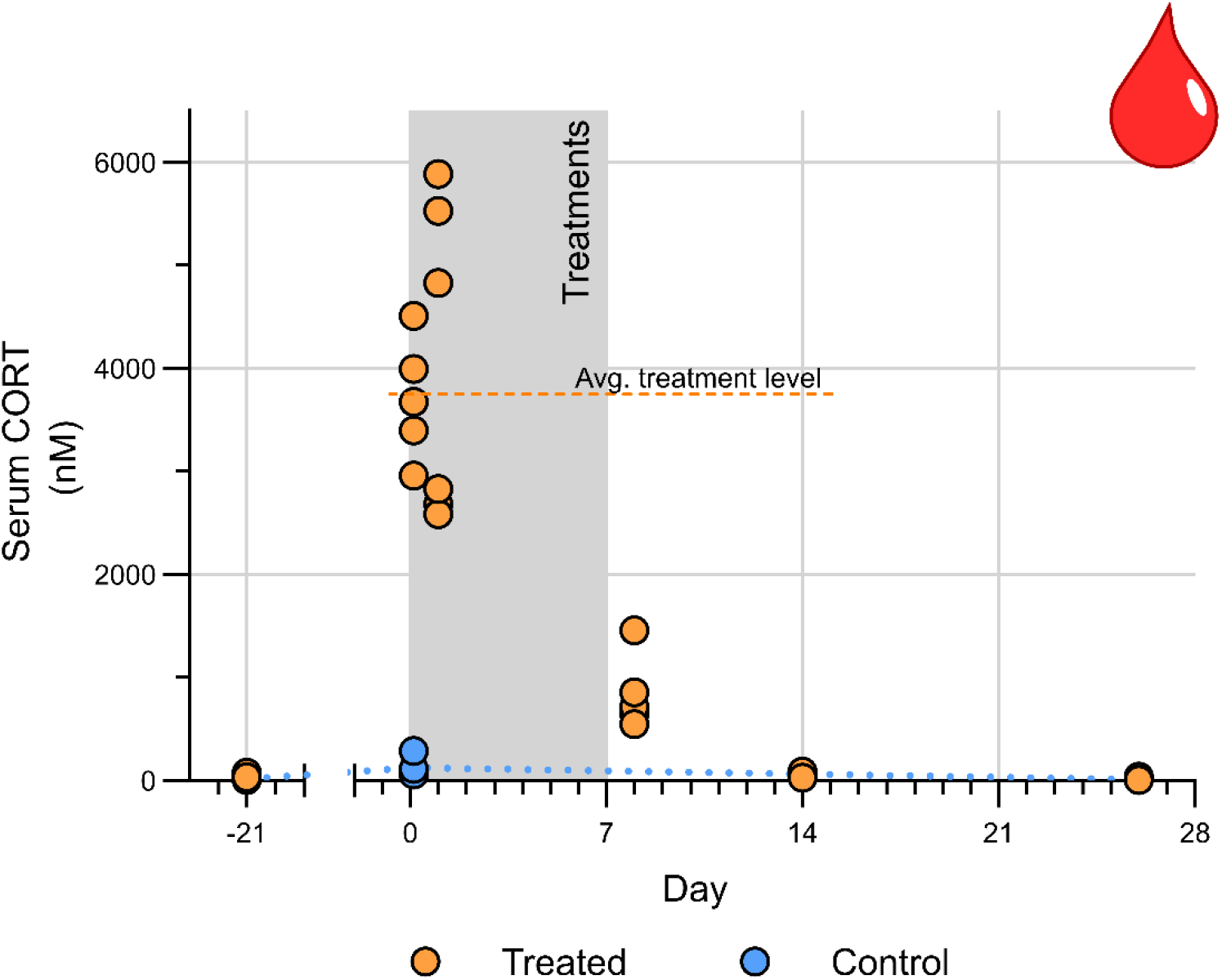
Time-concentration profile of serum CORT in ADX animals. The geometric mean of the serum concentrations in the treatment period for subjects injected with CORT has been included as a dashed line. The geometric means of the control animals have been connected with a dotted line.

The general time-concentration profile of CORT in hair was largely inconsistent with the trend predicted by the Classic model of hGC kinetics (Figure 5). A rapid increase in hair CORT levels was seen in relation to treatments, with the highest levels recorded on Day 8, *i.e*., 24 hours after treatments were discontinued (95 % CI: 377-3,970 ng/g, n = 5). The trend was consistent with what was found in Experiment 1. The levels were, however, lower than those measured on Day 7 of treatments in Experiment 1 (a time point that was not included), suggesting that actual peak levels during treatments may have been higher. When treatments were stopped, the hair CORT levels dropped markedly (Day 14 levels: 95 % CI: 422-858 ng/g, n = 5).

**Figure 5.**
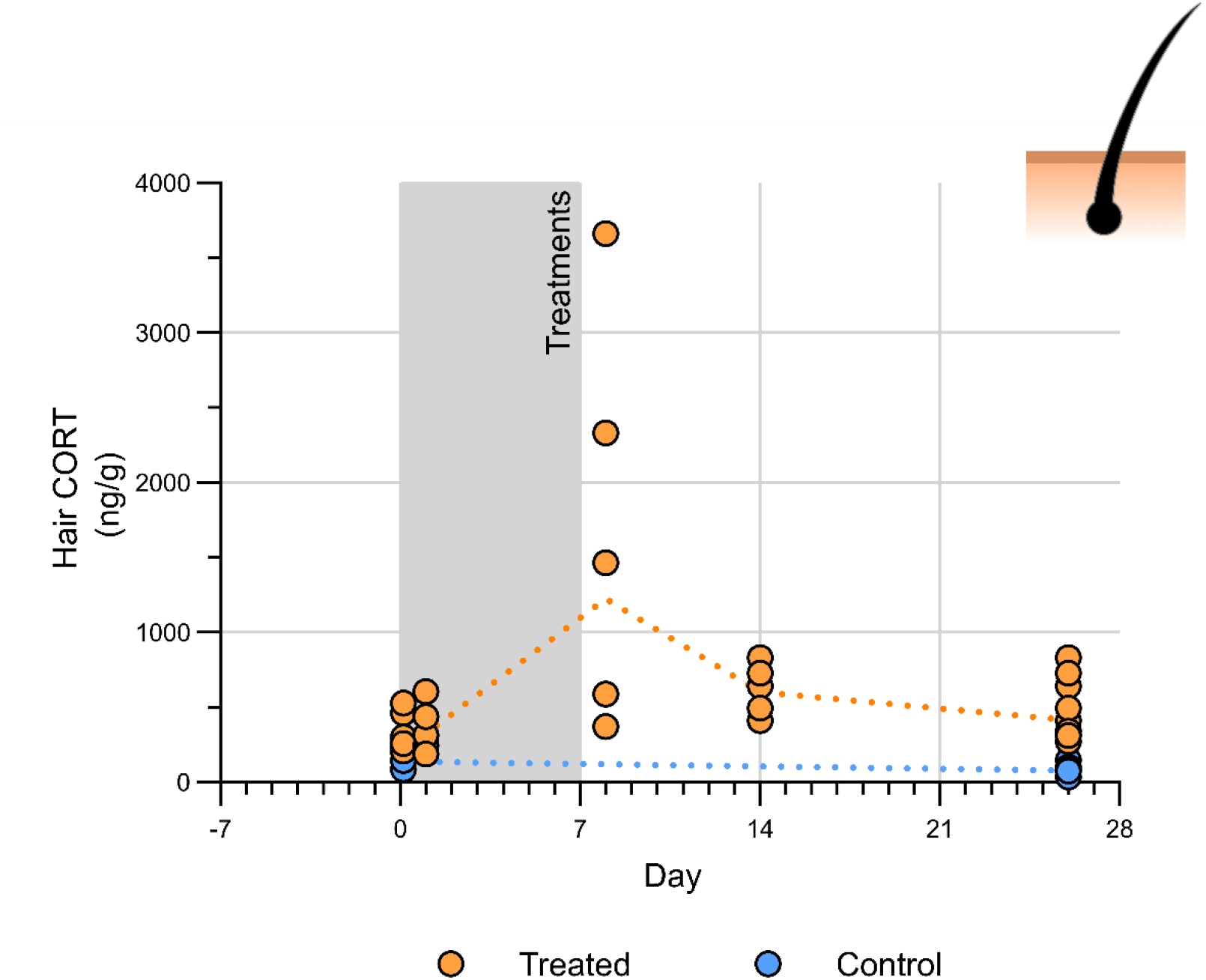
Time-concentration curves for serum and hair CORT in relation to one week’s treatments. The dotted lines connect the geometric means of the two groups of ADX animals.

Focusing on the modest hair CORT levels measured following the first treatment (3 and 24 hours after injection) and at the end of the study, two important findings were noted (Figure 6). Already after three hours, an elevation in hair CORT was seen in the treated animals, when compared to the untreated controls (Welch’s t-test: t_5.7_ = −2.96, p < 0.05). This elevation was found in a separate cohort of animals also 24 hours after the first injection (t_7.0_ = −2.95, p < 0.05).

On day 26, nineteen days after treatments were stopped, the hair CORT levels of the treated animals could still be distinguished from the adrenalectomized controls (t_14.0_ = −7.03, p < 0.001). Whereas this suggests that traces of a CORT response can be retained by hairs for a longer time, this elevation was of the same magnitude as that found three hours after the first injection (CORT treated animals sampled on Day 26 cannot be distinguished from those sampled at 3 hours: t_5.7_ = 0.42, p = 0.69).

**Figure 6.**
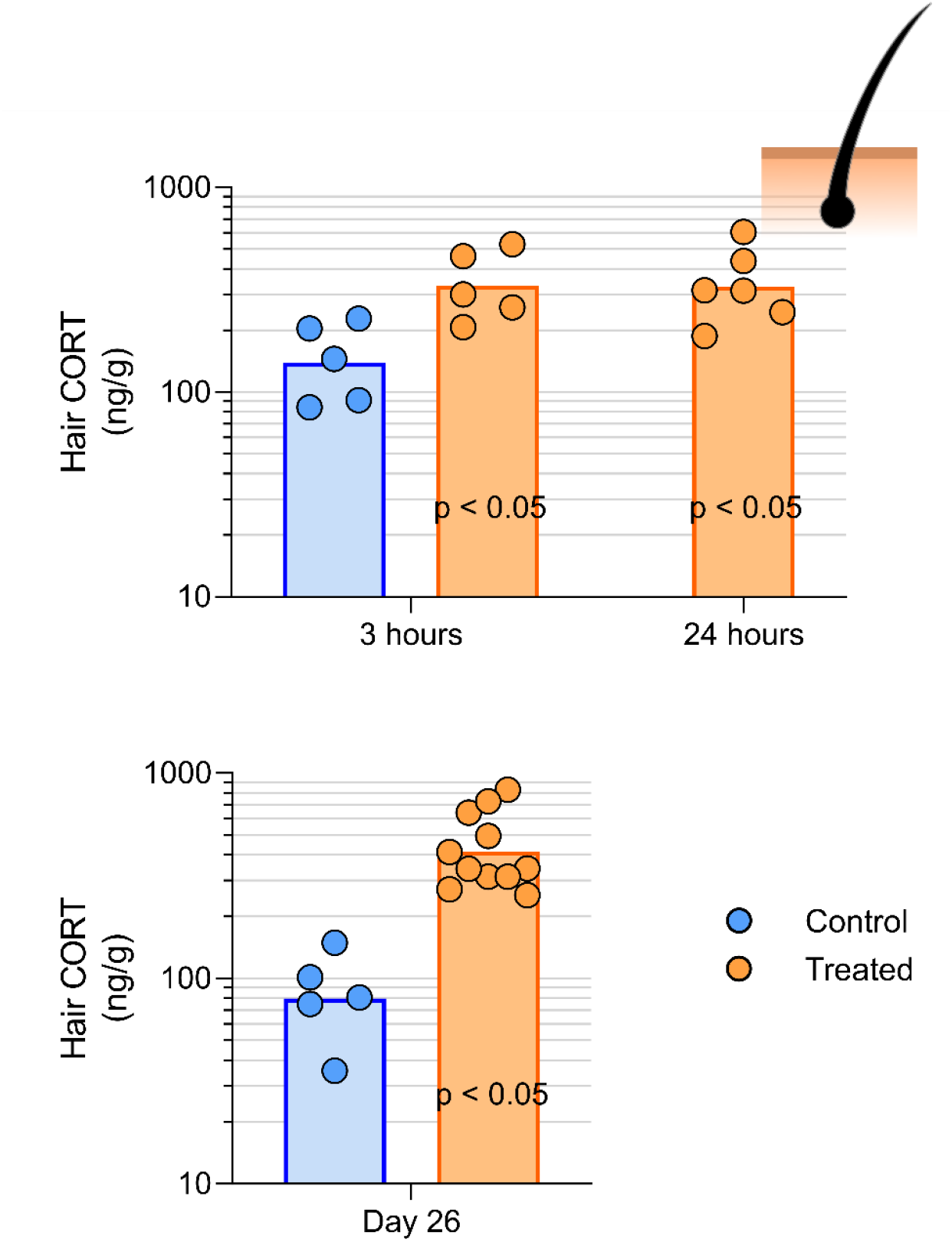
Closer inspection of hair CORT levels at 3 hours, 24 hours and Day 26 in ADX animals.

## 4. Discussion

Using a simple experimental design, devised to minimize ambiguity of interpretations, we were able to demonstrate that the Classic model used to interpret a majority of all studies employing hGC analyses has to be rejected. The lag time between (artificial) stressor onset and increased levels of CORT found in hairs was found to be less than three hours (and verified again at 24 hours); this is considerably faster than what is predicted by the Classic model. The highest hGC concentrations recorded in the present study were found on the last day of treatments (Day 7) in Experiment 1 (in animals which we had no reason to suspect had regrown adrenal tissue). This suggests that the hairs did indeed accumulate GCs over time. Finally, after treatments were stopped, hair CORT levels fell from Day 8 to Day 14 to Day 26. This demonstrates that GCs in hairs are not static, but diffuse in and out of the hair (or are eliminated). We had hypothesized that hair CORT levels would have returned to baseline levels within two weeks of the last treatment. This was not true, as trace levels of CORT were retained in hair samples collected on Day 26. Importantly, however, when comparing these levels to those found three hours after the first injection, we found the concentrations to be equivalent. The fact that a massive (synthetic) stressor lasting for eight days could not be distinguished from a stressor as recent as 3 hours in the past is strong evidence that the trace amounts of CORT retained in the hair cannot be used as a marker of past stress. It is worth noting that the elevated hair CORT levels found 3 hours, 24 hours, and 26 days after the first injection would probably be very hard to demonstrate in an intact animal. The ADX animals offer a completely noise-free template for measurements, an ideal scenario unlikely to resemble much of anything found outside of a laboratory.

In the end, the Alternate model describes experimentally obtained data fairly accurately. We will note that hairs retained CORT for a longer time than initially speculated, even though the resulting concentrations are unlikely to be usable in a context where CORT is considered a stress-sensitive biomarker. Nevertheless, a model where GCs diffuse into, along, and out of hairs describes the present evidence, and those previously published, best.

Where studies in the past have found results inconsistent with the Classic model, alternative explanations have been offered, suggesting that the Classic model is still, in essence, true. We will thus make a few observations pertaining to a few of these previously-highlighted explanations. Faster kinetics, with elevated hGC concentrations found shortly after a stressor (seen in, for example, Sharpley et al.^64^ or Cattet et al.^65^), have on occasion been attributed to surface contamination^65,66^. Whereas this cannot fully be ruled out in the present study, or indeed any other, we will note that we took steps to limit the effects of contamination. Rats do not possess sweat glands in close proximity to the sampled hair (rat sweat glands are only found on their paws)^67^ – ruling out contamination by sweat. This leaves two unchecked sources of externally deposited GCs – sebum and saliva (from grooming). Whereas the rats could also theoretically come in contact with urine or feces, it is unlikely to contaminate the hairs on their lower back (stereotypes aside, rats are fairly cleanly animals, and will separate latrine areas and sleeping sites in their cages if given the opportunity). The fur was visibly assessed to be clean when sampled. To limit the influence of external contamination, we employed the most common pre-treatment protocol^38,59^, washing our hair samples in isopropanol prior to sample extraction. This has been demonstrated to be effective in removing surface contaminants^68^ and is the protocol that is favored by a majority of groups utilizing hGC analyses^38^ (although other protocols do exist^69^). Where an induced stressor has not resulted in elevated hGC levels weeks after induction (seen in, for example, Ashley et al.^70^ or Crill et al.^71^) the magnitude of the stressor has been suggested to have been too small to elicit an elevation in hGCs. Where the stressor is sure to have been significant yet an elevation in hGCs cannot be found (*e.g*. the mothers in the study by Kapoor et al.^72^), where the amount of hGC has been found to be suspiciously low (*e.g*. in the study by Keckeis et al.^37^), or where it has disappeared faster than explained by the classic model (*e.g*. in, González-de-la-Vara et al.^73^) this has been ascribed abiotic factors such as rain^74^ and UV radiation from sunlight^75^. (In humans, other factors such as showering/shampooing have been discussed^76^.) In the present study we could be certain that the (artificial) stressor was sufficient to induce an elevation in hGC levels. Moreover, housed in shoebox cages, laboratory rats are not exposed to most of the theorized abiotic factors. Being a light-sensitive crepuscular species, lighting is kept low in the animals holding rooms and rats will favor sleeping in nests and under shelters. Consequently, of what little UV radiation fluorescent light tubes emit, laboratory rats are exposed to nearly none. Taken together, we find it extremely unlikely that our findings be explained through anything than the high diffusivity of CORT in hairs.

## 5. Conclusions

Hair glucocorticoid analyses have been used in studies that inform decisions that have consequences for human and animal health/welfare. It is thus disconcerting to find that the basis of their frame of interpretation has not been thoroughly investigated. There are numerous references made to GCs being incorporated into the growing hair, to a hair containing a record of the stress of its owner for its entire growth period, to a certain segment corresponding to a specific point in the past. These ideas are propagated through numerous narrative reviews, demonstrating the power of a good story. Long lists of convenient findings have been construed as evidence for the model, demonstrating the power of verification bias. Yet, when the evidence are summarized in systematic reviews^38,77^, little to no actual evidence has been found to suggest that hairs contain a record of past/historical stress. Evidence contrary to this have been presented however. Frankly, the findings of Kapoor and collaborators, that demonstrated that radiolabeled cortisol would migrate along the hair shaft^78^, should have been evidence enough for any laboratory utilizing hGC analyses to reconsider their interpretations. The Classic model proposed for hGC kinetics cannot hold true in the face of longitudinal movement of GCs.

In the present study we demonstrated the consequences of GC mobility. GCs will diffuse from the bloodstream into hairs, but they are not locked into place. They will diffuse throughout the hair and, as the concentration gradient changes, diffuse out of the hair again. With ADX rats, that have no central production of GCs, we find that traces of GCs remain in the hair for at least two weeks post a massive stressor (in the present study, an artificial one). But the trace amounts are too low to be usable as they can be overshadowed by a transient stressor. Given our results, we could speculate that a rat’s hair collects a usable record of experienced stressors in a time span of between a few hours and, perhaps, a couple of days in the past. The extent of the window is governed by the magnitude of the stressor, but also probably physiological characteristics such as the thickness of the hair. The exact kinetics would have to be determined on a per-species basis. The rat hair is a fair approximation, with respect to the thickness of the shaft, to the human (scalp) hair^79^. It is unlikely that any mammalian hair could contain a record of weeks, however, much less years. It should also be noted that hairs are more likely to retain corticosterone over time, when compared to cortisol, due to the former being more lipophilic^80^ (generally a characteristic recognized to be crucial for the retention of a molecule in hair^81^). The time from stressor onset to its appearance in the form of elevated GC content in hairs, and (importantly) the time for its disappearance as the stressor abates, needs to be established for a number of species. There are hundreds of published studies across numerous animal species that need to be reinterpreted with an updated model of hGC kinetics. Perhaps most pressing is the need to establish the hGC kinetics for humans.

We could, with our pilot investigation, demonstrate that an investigation similar to ours can be carried out also without adrenalectomies, provided a suitably controlled environment (we needed, however, to be certain, without a doubt, of our results – this level of certitude could not be established without adrenalectomized animals). All that is needed is daily treatments with high levels of GCs (cortisol or corticosterone, depending on species; ACTH challenges could alternatively be employed) and sampling of hair (and preferably blood) before, during, and after treatments using a shave-reshave protocol (summarized in Figure 1). This template enables investigators to easily determine the hGC kinetics of their species-of-study – whether voles, bears or humans. It also enables researcher of a skeptical bent to easily replicate our study, to verify for themselves that hairs do not contain a historical record of past stressors. Given the simple design, we, in fact, consider it highly likely that a similar study has been carried out in the past, but that the study results have been stowed away in a desk drawer as the researchers have found the results hard to interpret or believe (or hard to publish).

## Supporting information

Subject characteristics and drop-outs

Raw data (csv)

## 6. Acknowledgements

We are grateful of the expert technical assistance provided by Trine Marie Ahlman Glahder, Daniel Kylmann Hansen, and Helle Runchel Porsdal.

## 7. Financing and conflict of interest statement

This study was in part funded by the Center for Applied Laboratory Animal Research (CALAR) and the purchase of laboratory equipment was made possible through Brødrena Hartmanns Fond. The financers had no part in designing/interpreting the study and the authors have no conflicts of interest to report.

