## Supplementary material for "Hair glucocorticoids are not a historical marker of stress – exploring the time-scale of corticosterone incorporation into hairs in a rat model": Subject characteristics and drop-outs

**Supplemental methods: Subject characteristics and drop-outs**

Important for the study design was to exclude individuals who regenerated adrenal tissue, as endogenous glucocorticoids could potentially confound the results. Employing a better-safe-than-sorry approach we would rather exclude an animal too many than too few. Importantly, in order to not inadvertently bias the study, we chose, whenever possible, to not analyze hair samples from individuals suspected of adrenal regeneration.

Subjects were excluded based on weight-gain, hair growth rates, corticosterone concentrations in serum, and, ultimately, necropsies. Adrenalectomized (ADX) rats put on less weight than animals with functional adrenal tissue[1], their hair grows faster[2], and they will – obviously – have lower levels of corticosterone in serum (in the ideal situation, ADX rats will only have cross-reacting steroid molecules in circulation, but extra-adrenal production of corticosterone occurs[3]). Necropsies can reveal the presence of a regenerated adrenal[4], however, ectopic adrenal tissue can form following an adrenalectomy nearly anywhere[4,5]. Thus, a combination of observations, measurements and necropsies have to be used to determine individuals successfully depleted of adrenal tissue.

ADX rats consume saline rather than tap water to maintain their osmotic balance[6], yet cages with ADX animals were also provided with a bottle of tap water. Animals regenerating adrenal tissue will start consuming also tap water as they regain circulating aldosterone levels. This was used as an early detection system. Cages of ADX animals where tap water was lost in volumes that could not be explained as dripping from the bottle were identified as cages where one or multiple animals were likely to have regenerated adrenal tissue. We have omitted the cage-level water intake in the tables below, settling for reporting individual-level measurements.

Below, we have listed all ADX subjects from Experiment 1, some subject characteristics and, if they were excluded, the reason for doing so. Suspected adrenal regrowth was noted based on a reduced rate of hair-growth (post shaving), increased consumption of water (as opposed to saline), and a markedly increased weight-gain. A cut-off was set at 100 mM corticosterone in serum for separating ADX animals from animals who had (potentially) regenerated their adrenals. Regenerated adrenals were noted during necropsies by two blinded veterinarians. Due to risk of ectopic adrenal regrowth, necropsies were not alone considered reliable as a single criterion of exclusion. That is: all animals where a regenerated adrenal was found were excluded but, by contrast, an animal could still be excluded based on circulating levels of CORT while no adrenals could be found. Euthanized animals were necropsied, but the (attending) veterinarian did not specifically look for regenerated adrenals. In addition to the animals listed below, five animals died/were euthanized before they were assigned an ID. End of study blood samples were not collected from animals 1-1 through 1-9.

***ADX animals from Experiment 1.***

| Subject | Included/ excluded/ euthanized | End of study body weight (g) | Suspected adrenal regrowth | Baseline serum CORT (mM) | End of study serum CORT (mM) | Regenerated adrenal found |
|---|---|---|---|---|---|---|
| 1-1 | Included | 373 | No | 0.99 | - | No |
| 1-2 | Included | 410 | No | 1.57 | - | No |
| 1-3 | **Euthanized** | - | - | - | - | - |
| 1-4 | Included | 465 | No | 1.24 | - | No |
| 1-5 | Included | 367 | No | 1.10 | - | No |
| 1-6 | **Euthanized** | - | - | - | - | - |

| | | | | | | |
|---|---|---|---|---|---|---|
| 1-7 | Included | 464 | No | 8.21 | - | No |
| 1-8 | Included | 470 | No | 2.88 | - | No |
| 1-9 | **Euthanized** | - | - | - | - | - |
| 1-28 | **Excluded** | 529 | **Yes** | **329.68** | **325.79** | **Yes** |
| 1-29 | **Excluded** | 503 | **Yes** | 12.80 | **464.99** | No |
| 1-30 | **Excluded** | 512 | **Yes** | 18.06 | **159.75** | **Yes** |
| 1-31 | Included | 385 | No | 2.76 | 88.91 | No |
| 1-32 | **Excluded** | 666 | **Yes** | **367.33** | **433.74** | **Yes** |
| 1-33 | **Excluded** | 629 | **Yes** | 39.68 | **227.87** | **Yes** |
| 1-34 | **Euthanized** | - | - | - | - | - |
| 1-35 | Included | 464 | No | 6.40 | 96.04 | No |
| 1-36 | **Excluded** | 507 | **Yes** | **321.93** | **275.34** | No |
| 1-37 | Included | 406 | No | 5.13 | 2.51 | No |
| 1-38 | Included | 402 | No | 4.64 | 1.89 | No |
| 1-39 | Included | 436 | No | 7.63 | 11.46 | No |
| 1-40 | **Excluded** | 494 | No | 11.75 | **119.09** | No |

Below we have listed all animals (ADX throughout) from experiment 2, using the previously defined characteristics. Hair from excluded animals was not analyzed, with one exception: Animal 2-5 was excluded based on end of study serum CORT, which were analyzed in parallel with hair samples.

***Individuals from Experiment 2.***

| Subject | Included/ excluded/ euthanized | EoS body weight (g) | Suspected adrenal regrowth | Baseline serum CORT (mM) | End of study serum CORT (mM) | Regenerated adrenal found |
|---|---|---|---|---|---|---|
| 2-1 | **Excluded** | 567 | **Yes** | **374.63** | **> Max** | **Yes** |
| 2-2 | Included | 450 | No | 19.45 | 15.73 | No |
| 2-3 | Included | 515 | No | 7.40 | 21.15 | No |
| 2-4 | Included | 452 | No | 8.45 | 7.58 | No |
| 2-5 | **Excluded** | 590 | No | 84.95 | **113.28** | No |
| 2-6 | Included | 503 | No | 25.39 | 8.05 | No |
| 2-7 | Included | 470 | No | 57.26 | 6.95 | No |
| 2-8 | Included | 445 | No | 22.51 | 18.57 | No |
| 2-9 | Included | 475 | No | 24.76 | 1.73 | No |
| 2-10 | **Excluded** | 695 | **Yes** | **183.94** | **> Max** | **Yes** |
| 2-11 | Included | 540 | No | 20.70 | 23.21 | No |
| 2-12 | **Euthanized** | - | - | - | **-** | **-** |
| 2-13 | Included | 622 | No | 26.17 | 75.61 | No |
| 2-14 | Included | 460 | No | 12.09 | 9.01 | No |
| 2-15 | Included | 548 | No | 11.85 | 38.19 | No |
| 2-16 | Included | 492 | No | 30.01 | 3.66 | No |
| 2-17 | Included | 440 | No | 15.02 | 2.78 | No |
| 2-18 | Included | 453 | No | 77.97 | 16.88 | No |
| 2-19 | **Excluded** | 635 | **Yes** | **144.94** | **> Max** | **Yes** |
| 2-20 | Included | 455 | No | 22.55 | 26.07 | No |

| | | | | | | |
|---|---|---|---|---|---|---|
| 2-21 | Included | 463 | No | 16.27 | 22.76 | No |
| 2-22 | **Excluded** | 550 | **Yes** | **136.56** | **204.22** | **Yes** |
| 2-23 | **Excluded** | 526 | **Yes** | **> Max** | **91.70** | **Yes** |
| 2-24 | Included | 513 | No | 25.39 | 19.58 | No |
| 2-25 | **Excluded** | 600 | **Yes** | 32.47 | **> Max** | **Yes** |
| 2-26 | Included | 477 | No | 37.88 | 7.27 | No |
| 2-27 | Included | 483 | No | 57.15 | 15.54 | No |
| 2-28 | Included | 540 | No | 51.45 | 15.78 | No |
| 2-29 | Included | 460 | No | 19.42 | 3.00 | No |
| 2-30 | Included | 462 | No | 20.88 | 7.71 | No |
| 2-31 | Included | 460 | No | 65.28 | 7.70 | No |
| 2-32 | Included | 467 | No | 57.50 | 3.49 | No |
| 2-33 | Included | 463 | No | 29.25 | 2.05 | No |
| 2-34 | **Excluded** | 740 | **Yes** | 77.74 | **301.60** | **Yes** |
| 2-35 | Included | 465 | No | 30.53 | 5.32 | No |
| 2-36 | Included | 472 | No | 25.40 | 16.37 | No |
