## Supplementary material for "Hair glucocorticoids are not a historical marker of stress – exploring the time-scale of corticosterone incorporation into hairs in a rat model": Raw data (csv): Explanation for dataset.txt

Variables in dataset
----------------------------
Experiment
	1 or 2, as outlined in the associated manuscript

Subject ID
	Unblinded unique ID for each subject. 
	Note that subjects have been sampled more than once.

ADX
	Adrenalectomized individual, yes (1) or no (0).

Treatment
	Assignment to either treatment group (1) or control (0). 
	Note that this does not mean that the subject was treated at the time of sampling. 

Time
	Time point, in days, relative to start of treatments. 
	Note that 0 refers to the time point 3 hours after first treatment. 

Serum_CORT
	Concentration of corticosterone in serum. 
	Unit is nM (= nmol/l)

Hair_CORT
	Concentration of corticosterone estimated in hair.
	Unit is ng [corticosterone]/g [hair].

Excluded_data
	Indicates whether data was excluded (1) or retained (0) in the final analyses.
	Refer to specific supplemental materials document on why individuals were excluded from the experiments. 
